# A systematic genotype-phenotype map for missense variants in the human intellectual disability-associated gene *GDI1*

**DOI:** 10.1101/2021.10.06.463360

**Authors:** Rachel A. Silverstein, Song Sun, Marta Verby, Jochen Weile, Yingzhou Wu, Marinella Gebbia, Iosifina Fotiadou, Julia Kitaygorodsky, Frederick P. Roth

## Abstract

Next generation sequencing has become a common tool in the diagnosis of genetic diseases. However, for the vast majority of genetic variants that are discovered, a clinical interpretation is not available. Variant effect mapping allows the functional effects of many single amino acid variants to be characterized in parallel. Here, we combine multiplexed functional assays with machine learning to assess the effects of amino acid substitutions in the human intellectual disability-associated gene, *GDI1*. We show that the resulting variant effect map can be used to discriminate pathogenic from benign variants. Our variant effect map recovers known biochemical and structural features of *GDI1* and reveals additional aspects of *GDI1* function. We explore how our functional assays can aid in the interpretation of novel *GDI1* variants as they are discovered, and to re-classify previously observed variants of unknown significance.

## Background

Next-generation sequencing is now routinely practiced in the diagnosis of genetic conditions. However, the usefulness of these methods is limited by our ability to interpret the genetic variants that are discovered. The Genome Aggregation Database (gnomAD) (1), has amassed over 4.6 million unique missense variants present in the human population. Of these missense variants, 99% are rare (minor allele frequency < 0.5%) (2) and only 13% have a definitive clinical interpretation available on ClinVar (3). Therefore, methods to close the gap between variant identification and interpretation are needed.

Several approaches to variant interpretation are available, including genome wide association studies (GWAS), family segregation analysis, functional assays, and computational prediction of variant effects. Of these, GWAS and computational prediction can both be used to interpret data at a scale commensurate with the numbers of human genetic variants. However, GWAS is of limited value for the interpretation of rare variants due to limited statistical power and error in associations that is increased due to small sample sizes (4). Current computational prediction approaches are considered at best weak evidence for clinical variant interpretation (5). Functional assays have traditionally been used to test variants on an individual basis, but these experiments are resource-intensive and this evidence is unlikely to be available at the time a newly-discovered variant is first classified. However, it has become possible to perform multiplexed assays of variant effect (MAVE), enabling the testing of functional effects for large numbers of missense variants in parallel (2,6–8). For example, a framework for variant effect mapping of human genes by complementation in *S. cerevisiae* has been previously described and applied to multiple genes (8–10). This framework has been shown to identify, at stringent confidence thresholds (90% precision), two to three times more pathogenic variants than are identified by computational prediction alone (8–10). Here, we apply this framework to carry out large-scale testing of missense variants of human *GDI1*, one of multiple genes on the X chromosome that have been found to contain mutations causing X-linked non-syndromic intellectual disability (11).

The *GDI1* gene encodes the protein GDI1 (Rab GDP dissociation inhibitor alpha). In mammals, GDI1 is expressed primarily in the brain and is necessary for the control of endocytic and exocytic pathways in neurons and astrocytes through the spatial and temporal control of numerous Rab proteins (12,13). GDI1 functions to extract inactive GDP-bound Rab from membranes by binding and solubilizing the genranylgeranyl anchor (a post-translational modification at C-terminal cysteine residues which anchors Rabs to membranes) (14). *GDI1*-null mouse models show deficits in short- and long-term synaptic plasticity and behavioral phenotypes including alteration of hippocampus-dependent forms of short-term memory, spatial working memory and associative fear-related memory (12). In humans, *GDI1* loss-of-function variants can cause non-syndromic intellectual disability (ID), characterized by cognitive impairment in the absence of other symptoms or physical anomalies (11). The form of ID caused by *GDI1* variants follows an X-linked semi-dominant pattern of inheritance, with hemizygous males being most severely affected and female carriers showing milder or no symptoms (15,16).

As a common condition which has been estimated to affect up to 3% of the general population (11), ID presents a diagnostic challenge due to its many potential causes. Alterations in over 700 genes have been associated with ID, few of which are frequently-occurring (17,18). Separating causal from benign genetic variation in ID patients is therefore a significant clinical challenge. Indeed, although an etiological diagnosis brings substantial benefits for patients and their families (19), including more accurate prognosis, genetic counselling on recurrence risk, and earlier access to resources within the community and specialized education programs, only ∼30% of ID patients receive an etiological diagnosis (20,21). Proactive functional testing for variants in genes associated with ID could aid in the identification of causal variants and facilitate earlier etiological diagnosis.

Here, we present large-scale measurements of the functional effects of missense variation in *GDI1*. Variant assay results are consistent with our knowledge of GDI1 function. A comparison of variant scores with ClinVar annotations suggests that the map will prove useful in assigning pathogenicity to genetic variation.

## Results

### Multiplexed yeast complementation efficiently identifies damaging *GDI1* variants

To efficiently test the deleteriousness of *GDI1* missense variants, we used a previously-validated humanized yeast model system(22). In this system, the *Homo sapiens GDI1* (*HsGDI1*) can complement a temperature sensitive allele of the orthologous *Saccharomyces cerevisiae* gene Gdi1 (*Sc*Gdi1 (Ts)) and thereby restore yeast growth at restrictive temperatures. Importantly, pathogenic variants of *HsGDI1* (L92P and R423P) showed a reduced ability to complement *Sc*Gdi1(Ts) (22). This supported the possibility of a yeast-based functional assay of *HsGDI1* variants, which we scaled up in order to test large numbers of missense variants in parallel (fig. 1a).

**Figure 1:**
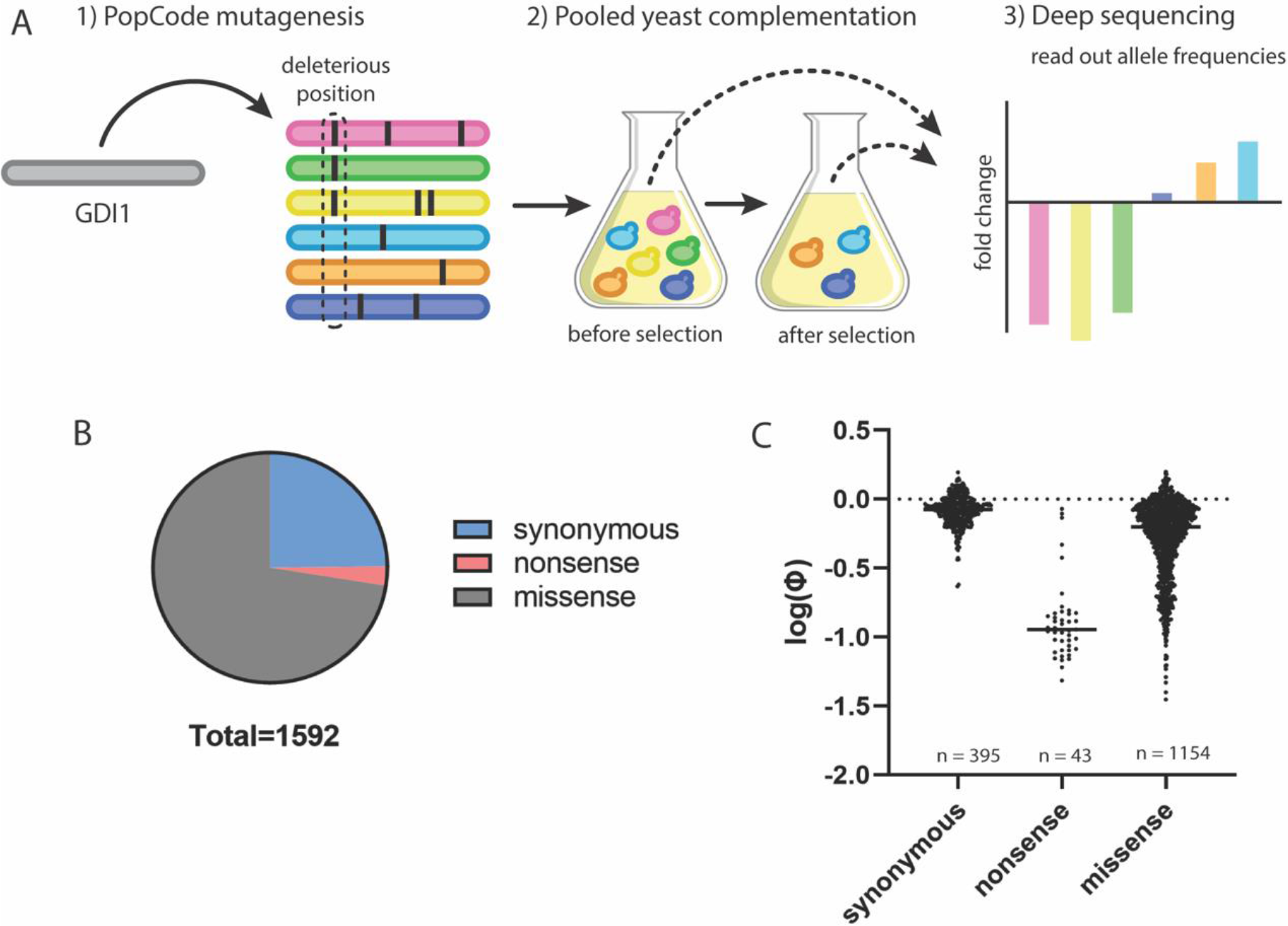
High throughput yeast complementation screen separates synonymous and nonsense *GDI1* variants. a) Graphical overview of the variant effect mapping framework. b) Number of well-measured variants recovered from the complementation screen. c) Log(ϕ) values comparing pre- and post-selection variant frequencies for all well measured synonymous, nonsense and missense *GDI1* variants.

Mutagenesis of the *HsGDI1* open reading frame (ORF) was performed using a previously-described pooled mutagenesis approach, Precision Oligo-Pool based Code Alteration or “POPCode” (8), which uses oligonucleotide-directed codon randomization to yield a library of single-codon *GDI1* variants. Following mutagenesis, the variant library was cloned into yeast expression vectors and transformed *en masse* into a *S. cerevisiae* strain carrying the temperature sensitive *Sc*Gdi1(Ts) allele. The yeast library was then grown competitively at restrictive temperatures to induce selection for cells containing functional *HsGDI1* variants.

The library of *HsGDI1* ORFs was extracted from both pre- and post-selection yeast populations, and sequenced deeply (with each position being observed in ∼2 million reads). The deep sequencing approach used was TileSeq (8), involving amplification and paired-end sequencing of 12 “tiles”, each ∼100 nucleotides in length, that together cover the length of the *GDI1* ORF. In order to decrease the rate of variants called erroneously due to sequencing error, only variants that were detected in both forward and reverse reads were accepted. In total, 5534 unique amino acid changes were detected. To understand the rate at which missense variants are detected due to PCR or sequencing errors, we also sequenced a ‘mock library’ derived from a wild-type clone. These data were used to filter out variants that were not represented at high enough frequencies in the pre- or post-selective pools to rule out the possibility that they were detected due to PCR and sequencing error alone (see materials and methods). Even after this filter, variants that were present at lower frequencies in the pre-selection library showed poorer agreement between replicates (fig. S1) and poorer correlation with PROVEAN (23) scores (fig. S2b). We therefore identified a set of high confidence variants by further removing variants that had been detected at a frequency lower than 2×10^−4^ in the original library. After filtering, 1730 high confidence variants remained, covering 1154 unique amino acid changes (19% of all possible amino acid substitutions and 45% of possible amino acid substitutions accessible through alteration of a single nucleotide (fig. 1b).

For each variant, the ratio (ϕ) of frequency in post-to pre-selective pools was used to infer variant functionality. Indeed, we saw a distinct separation between log(ϕ) values for synonymous variants, which would generally be expected to fully complement the *Sc*Gdi1(Ts) allele, and log(ϕ) values for stop codon variants, which would generally be expected to completely fail to complement (fig. 1c). Most missense variants appeared wild-type-like in their ability to complement, some were null-like, and many had intermediate effects (fig. 1c).

### A variant effect map for *GDI1*

Log(ϕ) values were rescaled to define a “fitness score” for each variant, representing the ability of that variant to complement the *Sc*Gdi1(Ts) allele (see materials and methods). With the goal that a fitness score of 1 represents a fully-functional protein and a fitness score of 0 represents complete loss of function, we rescaled log-ratios such that the median log ratio of synonymous variants was 1 and the median log ratio of variants containing a premature stop codon was 0 (medians shown in fig. 1c). When calculating median log ratios, we included only high confidence measurements (SD < 0.3) and, because nonsense mutations near the C-terminus result in less severe loss of complementation, we only considered nonsense mutations within the first 400 amino acids of the *GDI1* ORF (fig. S3). In order to estimate fitness scores for the remaining 80% of amino acid changes and refine scores of variants that were less well measured, we applied a previously-described imputation pipeline (24). This pipeline uses the Gradient Boosted Tree method to impute missing values based on intrinsic features of the data set including average fitness of nearby variants, amino acid substitution matrix scores (BLOSUM100 (25)), and variant effect scores predicted by computational methods including PolyPhen-2 (26), and PROVEAN (23). To avoid low-confidence predictions based on limited experimental data, imputation was not performed for amino acid positions with fewer than 3 well-measured variants. The result was a ‘variant effect map’ encompassing the majority of all possible amino acid substitutions in *GDI1* (fig. 2). The most important features for predicting fitness scores in this data set were the average fitness scores of the three most similar variants at the same amino acid position, followed by BLOSUM100, PolyPhen2, and PROVEAN scores (fig. S4).

**Figure 2:**
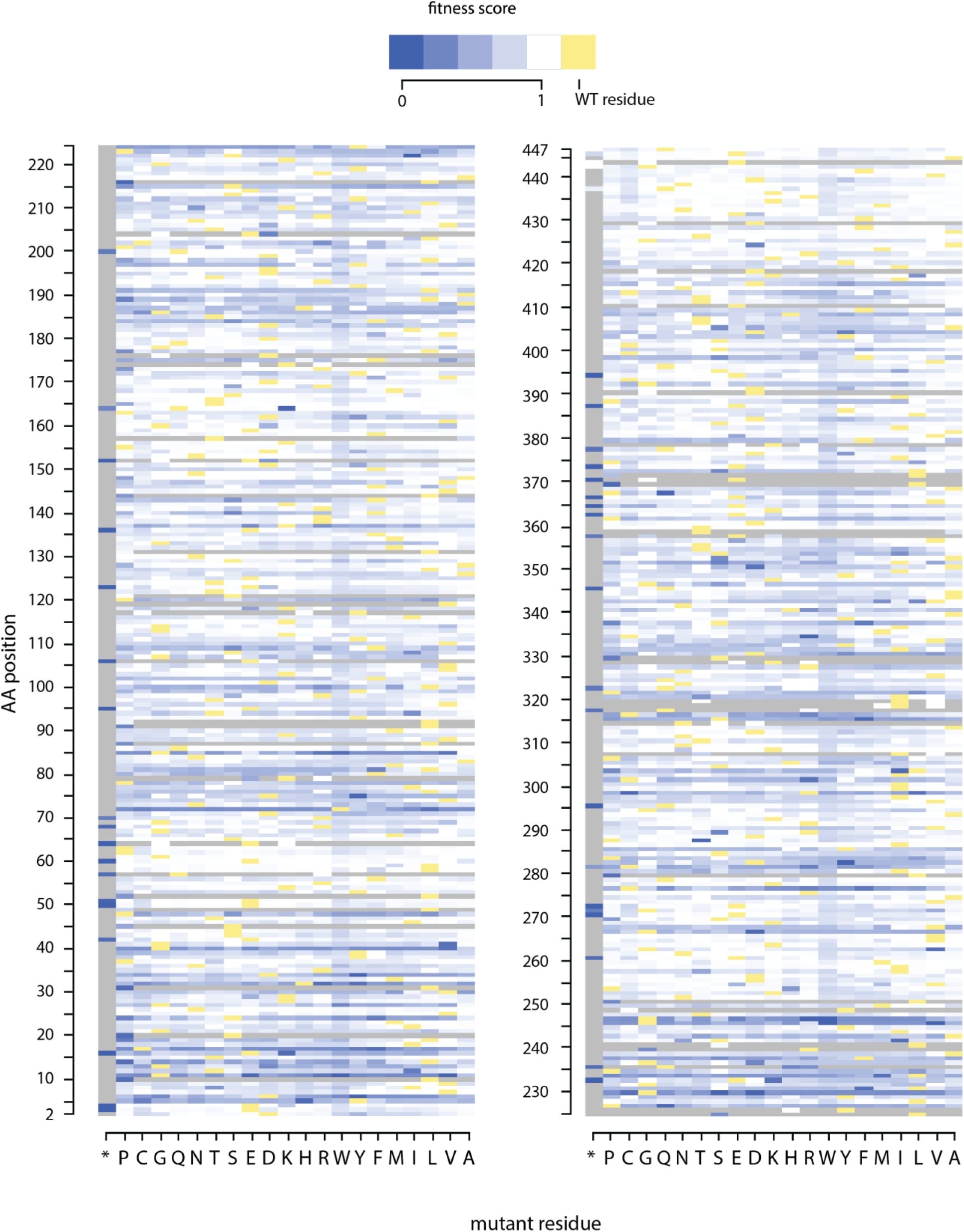
*GDI1* variant effect map. A *GDI1* missense variant effect map resulting from the complementation screen coupled with imputation and refinement by machine learning. Fitness scores of 0 (blue) represent the median behavior of complete loss of function variants (based on observed fitness of nonsense variants) and fitness scores of 1 (white) represent wild type-like function (based on observed fitness of synonymous variants). Yellow tiles represent the wild type amino acid at that position. Gray tiles represent substitutions for which scores were not imputed due to insufficient data for substitutions at that amino acid position.

### Our variant effect map is consistent with known biochemical features of GDI1

The GDI1 protein contains four sequence conserved regions (SCRs), SCR1, SCR2, SCR3A and SCR3B, common to all members of the Rab-GDI/CHM superfamily (27). Together, SCR1 and SCR3B form a Rab-binding platform at the apex of the GDI1 structure (27,28) (fig. 3a). SCR3A contains a mobile effector loop (MEL) which constitutes a membrane receptor binding site as well as a helix flanking the lipid binding pocket (29,30). At its N-terminal end, SCR2 contains the C-terminus–binding region (CBR), which forms an essential interaction with the C-terminus of Rab (28).

**Figure 3:**
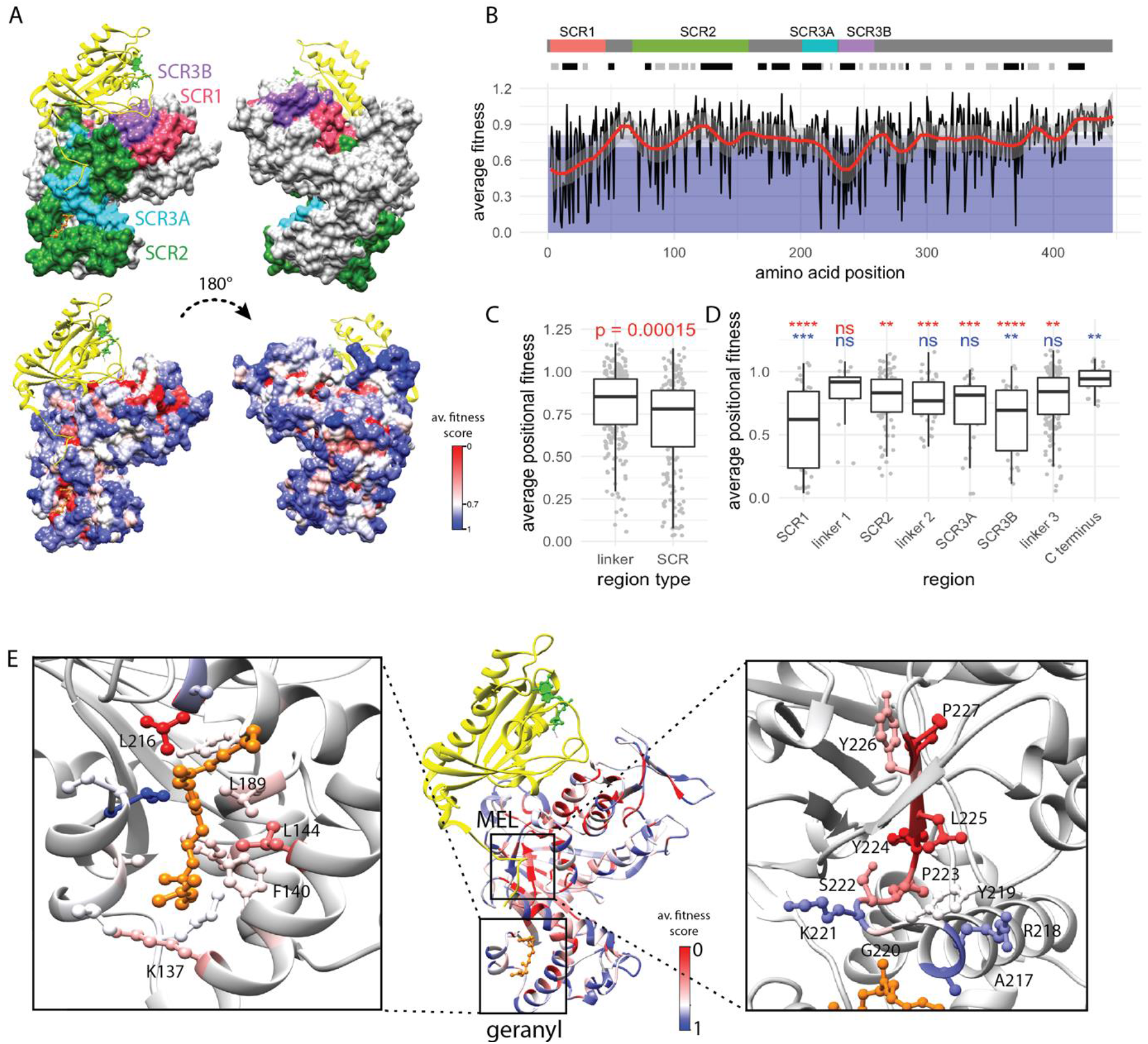
Fitness scores enable structure-function analysis of *GDI1*. (a) Homology model of human GDI1 (colored surface) modeled on the structure of *S. cerevisiae* RabGDP-dissociation inhibitor in complex with prenylated YPT1 GTPase (yellow ribbon). In the bottom panel, residues are colored according to their average positional fitness scores with 0 representing null-like scores (red) and 1 representing wild type-like scores (blue). (b) Average fitness score of all variants at each amino acid position (black line) overlaid with a smoothed summary curve (red). The dark blue region of the plot represents fitness scores less than 0.72 (over 10 times more likely to be damaging than tolerated) and the white region represents fitness scores over 0.81 (over 10 times more likely to be tolerated than damaging). [Tolerated:damaging odds ratios were calculated as described in methods]. The tracks above the plot represent: depiction of GDI1 with sequence conserved regions (SCRs) common to all members of the GDI/CHM superfamily (top track) and; the secondary structures of human GDI1 as predicted by PSIPRED 4.0 (44) (bottom track; black = helix, gray = strand). (c) Average fitness scores of amino acid positions within non-conserved or “linker” regions versus sequence conserved regions. Significance level was determined using Wilcoxon signed-rank test. (d) Region-wise comparison of average positional fitness scores. Wilcoxon signed-rank tests were performed comparing each region to the “C-terminus” region (red asterisks) and to SCR2 (blue asterisks). Significance levels are denoted by: * (p<0.05), ** (p<0.01), *** (p<0.001), and **** (p<0.0001). e) Center: Ribbon representation of human GDI1 modeled on the structure of *S. cerevisiae* RabGDP-dissociation inhibitor in complex with prenylated YPT1 GTPase (yellow). GDI□ residues are colored by average positional fitness score. Left: side chains of all hydrophobic residues within 5A of the geranylgeranyl group (orange). Right: side chains of residues comprising the mobile effector loop and proximal beta strand.

To determine overall patterns of variant deleteriousness within GDI1, we took the average fitness score of all variants at a given amino acid position resulting in a “positional fitness score” (fig. 3b). As expected, average fitness was significantly lower in the sequence-conserved regions than in other parts of the protein (fig. 3c, 3d), supporting the notion that these regions are important for biological function. We modeled the sequence of *H. sapiens* GDI1 on the crystal structure of *S. cerevisiae* RabGDP-dissociation inhibitor in complex with prenylated YPT1 GTPase (28) (the yeast homolog of human Rab-1A). The conserved face of GDI1 constituting the Rab binding platform contains the majority of residues with low positional fitness scores (fig. 3a). Mutations in the SCR1 and SCR3B segments exhibited the lowest positional fitness on average (fig 3d), consistent with previous mutational analysis showing that disrupting these regions leads to decreased Rab binding and inability of GDI1 to extract Rab from membranes (27). Since the C-terminal non-conserved region showed a striking increase in average fitness scores around residue 425 (fig. 3b), we divided this region into two separate sections, “linker 3”, consisting of residues 460-424, and “C-terminus”, consisting of residues 425-447. Mutations in the 22 “C-terminus” residues were significantly less deleterious than those in linker 3 (Wilcoxon p<0.01). The non-conserved region between SCR1 and SCR2 (termed “linker 1”) also exhibited high fitness scores, suggesting that variation here is also well tolerated (fig. 3d).

Compared to SCR1 and SCR3B, variants in SCR2 were significantly less deleterious (Wilcoxon p<0.01, and p<0.001 respectively). On average, fitness scores of variants in SCR2 were comparable to those in the non-conserved region between SCR2 and SCR3A (termed “linker 2”) and the non-conserved linker 3 region (fig. 3d). Within SCR2, variants with the most severe fitness effects tend towards the N-terminal CBR segment (fig. 3b). However, altering any one of several hydrophobic residues within the helices flanking the lipid binding pocket, especially Leu216 and Leu144, also yielded low positional fitness (fig. 3e). The location of these residues, coupled with their average positional fitness scores, suggests that they may play an important role in geranylgeranyl binding.

Deleterious mutations within the SCR3A region were observed predominantly towards the C-terminus. Residues within the MEL region had moderate average positional fitness scores between 0.5 to 0.75. It was previously reported that when MEL mutations Arg218Ala, Tyr219Ala, and Ser222Ala are introduced into the corresponding positions of the yeast protein *Sc*Gdi1, they do not cause visible growth defects in yeast. However, when any one of these is introduced in combination, they can exacerbate the effects of partial loss-of-function variants elsewhere in *GDI1* (29). Our results show that single mutants Arg218Ala, Tyr219Ala, and Ser222Ala each result in modest loss of function with fitness scores of 0.75 +/- 0.18, 0.67 +/- 0.22, and 0.66 +/- 0.13 respectively (regularized standard error for fitness scores was calculated as described in materials and methods). It is possible that our competition-based assay was more sensitive to minor growth changes and thus able to detect growth defects not detected by spotting assays. While the study by Luan et al. only tested mutations in residues 218 - 222, we observed some variants just outside of this region to be extremely deleterious, especially a short β strand (termed β-strand e3 in Luan *et al*.) from residues Ser222 to Pro227 (fig. 3e). Despite the importance of this segment indicated by our map, a biological function for this strand segment has not been described.

### Relating fitness score to severity of intellectual disability

Severity of ID is highly variable with cases ranging from mild to profound (31). Although the severity of ID has been reported for only three *GDI1* missense variants have been reported to date, we explored whether there was potential for variant fitness scores to predict the severity of the associated ID phenotype. Males from a family with the Leu92Pro variant, were reported to suffer from mild to moderate ID (11,32). For this variant, we obtained a fitness score of 0.74 +/- 0.03. Individuals in a family carrying the Gly237Val variant were reported to have moderate ID (33), and we observed a corresponding lower fitness score of 0.55 +/- 0.07 for Gly237Val. Thus, the order of the fitness scores for these two variants agreed with the reported order of ID severity. We note however that, like 20 (80%) of the 25 variants listed in the ClinVar database, both Leu92Pro and Gly237Val are currently annotated as a variants of uncertain significance, highlighting the need for better tools for interpretation. Finally, a family carrying the Arg423Pro variant suffered moderate to severe ID (15). We did not observe Arg423Pro in our assay, and were only able to impute a score with necessarily higher estimated uncertainty (0.64 +/- 0.24). Although fitness scores may be predictive of ID severity; it is currently insufficient to draw this conclusion from only reported ID severity data.

### *GDI1* variant effect map predicts pathogenic variants with higher precision than computational methods alone

In order to test whether fitness scores from the *GDI1* map can provide useful evidence for determining variant pathogenicity, we wished to determine whether our variant effect map can be used to separate known benign from damaging alleles. Our set of presumed-damaging variants included the only variant currently annotated as pathogenic (Arg423Pro (15)) and the additional missense *GDI1* variants discussed above: Leu92Pro (11,32) and Gly237Val (33) based on evidence from clinical reports. Because the number of currently known human pathogenic variants is small, we also included four missense variants in highly-conserved regions which have been previously shown to inhibit the ability of GDI1 to extract Rab3A from membranes in rat synaptosomes, Tyr39Val, Glu233Ser, Met250Tyr, and Thr248Pro (27). To establish a reference set of presumed-tolerated variants, we extracted all variants in gnomAD that had been observed in male subjects (who are hemizygous for *GDI1* and less likely to be ID given that gnomAD excludes subjects with early-childhood disease). Although we cannot rule out the possibility that our set of presumed-damaging variants contains some tolerated variants, nor that our set of tolerated variants contains some damaging variants, we reasoned these sets would enable a conservative estimate of the ability of our scores to distinguish damaging from tolerated variation.

We observed that our sets of presumed tolerated and damaging alleles were well-separated based on fitness score (fig 4a). Although fitness scores for presumed-damaging variants showed a strong tendency to have lower scores, the lowest score amongst these was 0.5 and none were null-like. We next calculated a precision-recall curve (fig. 4b) showing, as we change the fitness score threshold below which a variant is deemed “damaging,” the trade-off between precision (fraction of below-threshold variants that are damaging) and recall (fraction of damaging variants that are below the threshold). For comparison we also provide precision-recall curves for commonly used computational predictors of variant effect including PolyPhen-2 (34), PROVEAN (23), and VARITY (35) (fig. 4b). Our variant effect mapping framework was able to identify 6 out of 7 damaging variants (87% recall) with 100% precision using a fitness score threshold of <0.68. We identified all damaging variants (100% recall) with 88% precision when a threshold fitness score of <0.74 was used. The most accurate computational predictors were VARITY and PolyPhen-2, which were each able to identify ∼75% of damaging variants with ∼75% precision.

**Figure 4:**
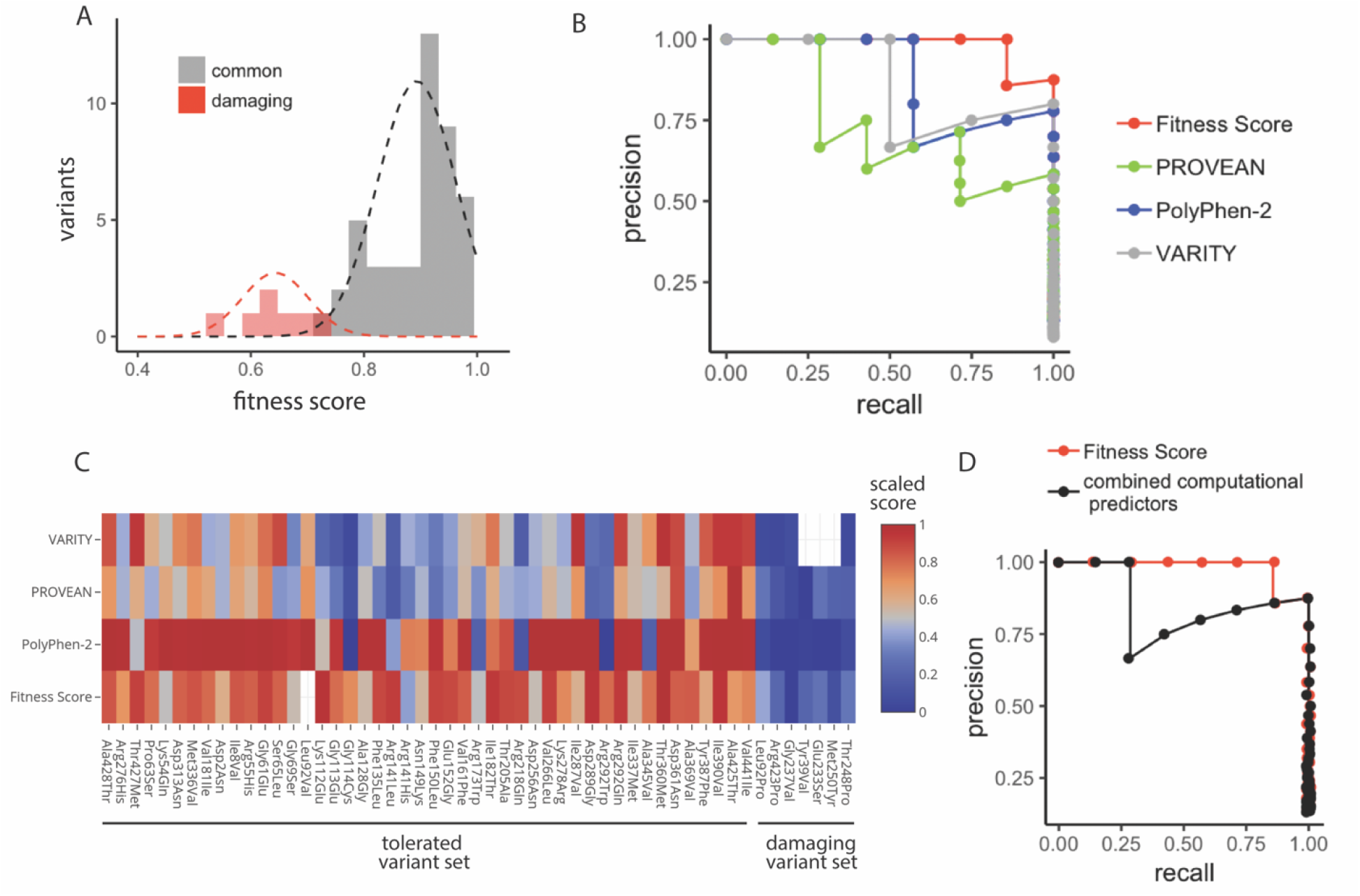
*GDI1* variant effect map separates damaging and common variants with higher precision than current computational methods. (a) Distribution of fitness scores for known damaging and known common (presumed tolerated) *GDI1* missense variants. Common variants are comprised of 46 missense variants listed in gnomAD which have been observed in at least one hemizygous individual. (b) Precision-recall curve for our fitness scores compared to various computational methods for variant interpretation. A sliding threshold was used for each score type starting at the lowest score; variants below this threshold were called as damaging. For each threshold value, the number of true damaging variants identified (true positives) and the number of benign variants identified in error (false positives) was evaluated. The precision [true positives/(true positives + false positives)] versus the recall [true positives/(true positives + false negatives)] is shown for each threshold value. c) Scaled fitness scores and computational predictor scores for all variants from our tolerated and damaging variant sets. Scores were scaled such that all score types range from 0 to 1 with 0 representing most damaging and 1 representing most tolerated. d) Precision recall curves for our fitness scores and for a “combined computational predictor score” which is the median of scaled PolyPhen-2, PROVEAN, and VARITY scores (scaling was performed as described in c).

Because single computational predictors are rarely used in isolation, we wondered whether a combined computational prediction score, encompassing data from PolyPhen-2, PROVEAN, and VARITY could separate damaging and tolerated variants with accuracy similar to our variant effect mapping framework. We rationalized that agreement between multiple computational prediction methods might be interpreted as stronger evidence for variant effect than a single computational method alone. We therefore scaled the PolyPhen-2, PROVEAN, and VARITY scores, and our fitness scores for the tolerated and damaging variant sets described above such that scores ranged from 0 to 1 (with 0 representing most damaging and 1 representing most tolerated). This allowed us to make comparisons between the different score types (fig. 4c). Notably, all the prediction methods were able to accurately identify all of the damaging variants (fig 4c). (Note that VARITY only generates predictions for single nucleotide variants, so scores were not generated for 3 out of 7 damaging variants). However, it can be seen that the increased accuracy of our VE mapping framework is due to the lack of false positives (prediction of a damaging variant/low fitness score when the variant is in fact tolerated). Moreover, the three computational methods tended to agree on many of the false positives, each assigning them low scores when the variant was in fact tolerated. For instance, Gly114Cys, Arg141Leu, Arg218Gln, and Arg292Trp were four particularly prominent false positives where each of the computational methods predicted a low score, however, the fitness score generated by our VE mapping framework correctly indicated that the variant was tolerated. Thus, we conclude that agreement between computational predictors does not necessarily appear to be an indicator of accuracy. To further illustrate this, wished to generate a “combined computational predictor score” which takes into account the agreement between different computational predictors. We therefore took the median of the scaled scores from the three computational prediction tools, reasoning that this would eliminate outliers where an individual prediction tool did not agree with the other two. When we plotted the precision recall curve for this “combined computational predictor” we found that it did not perform better than the individual predictors (fig. 4d). This is consistent with our notion that the shortcomings of the computational prediction methods are not due to individual outlying predictions.

### Using our VE map to interpret clinically-relevant missense variants

To facilitate the use of fitness scores as evidence to classify variants, we wished to calculate likelihood ratios that convey the extent to which one should raise or lower the probability that a variant is damaging, based on the fitness score. To this end, we estimated probability density functions that describe the distributions of scores from our presumed-damaging and -tolerated variant sets (see Methods). Then, the ratios of probability density for damaging and tolerated variants can be used to obtain a damaging:tolerated likelihood ratio for variants with any given fitness score. By this method, we determined that variants with fitness scores below 0.72 were over 10 times more likely to be damaging than tolerated and variants with fitness scores above 0.81 were over 10 times more likely to be tolerated than damaging.

We wondered whether our map could aid in the interpretation of GDI1 variants of unknown significance which have been observed in the clinic (fig. 5). The ClinVar database lists 25 missense variants in GDI1, only four of which currently has a definitive clinical interpretation. For 15 out of the 21 variants without a definitive interpretation, we were able to generate an interpretation of either “deleterious” or “tolerated” with odds ratios greater than 1:10 based on our VE map (fig. 5). In order to be conservative with our interpretations, any variants which had intermediate fitness scores leading to odds ratios less than 1:10 were labeled as “unknown”. Of note, we discovered four additional variants (in addition to those previously included in our “likely damaging” variant set) that, were found by our assay to be highly deleterious: R35W, G40V, R290S, and V381E. The latter three of these variants had almost null-like scores. This highlights the possibility that ID due to *GDI1* mutations is under-diagnosed due to current limitations in clinical variant interpretation.

**Figure 5:**
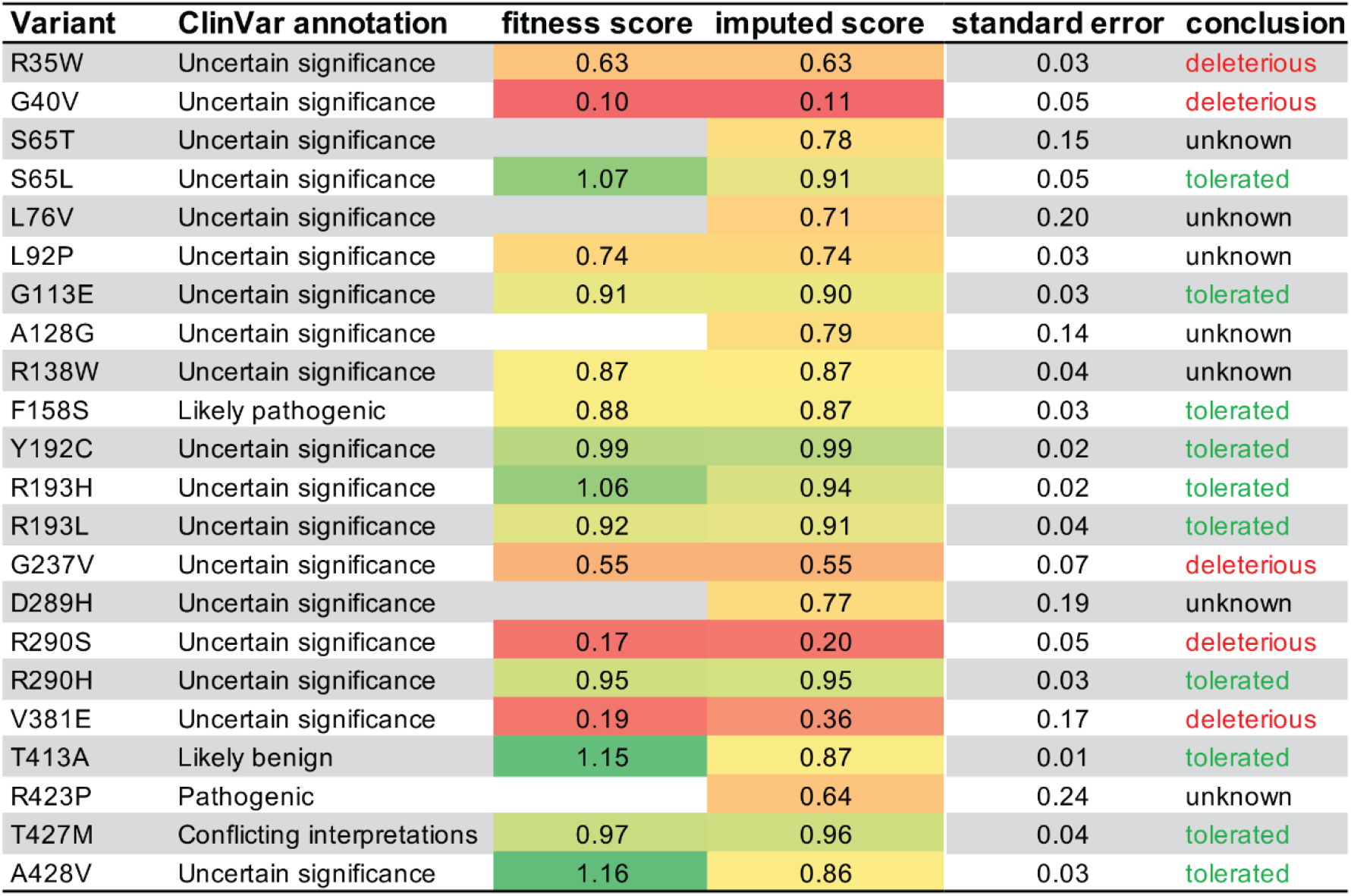
Interpretation of clinically-relevant GDI1 missense variants from the ClinVar database. Fitness scores (experimentally measured where available, and computationally imputed for all variants) are listed for all GDI1 missense variants listed on ClinVar. We concluded that a variant is “deleterious” where the damaging:tolerated odds ratio was greater than 1:10 and vice versa for “tolerated” variants.

Interestingly, Phe158Ser, annotated on ClinVar as “likely pathogenic” based on the amino acid change being located within in a conserved region (SCR2), was non-conservative with respect to amino acid properties, and was not observed as a common variant in the NHLBI Exome Sequencing Project (37). However, our map score for Phe158Ser ([0.875 +/- 0.03] originally, [0.870 +/- 0.03] post-refinement) does not provide strong evidence that this variant is damaging. Using our current model based on the distributions of known pathogenic and benign variants, and using no prior assumptions about the pathogenicity of Phe158Ser, a fitness score of 0.87 indicates the odds that the variant is damaging is less than 1:100. If our likelihood ratio calibration is accurate, then even given a very strong prior belief (P = 0.99) that this variant is damaging, the posterior odds would be less than 1:10.

## Discussion

Towards clinical variant interpretation, the likelihood ratios that we derived for each variant from our map could be discretized as strong, moderate, or supporting evidence for the functional impact of a variant, and combined with other evidence using American College of Medical Genetics and Genomics/Association for Molecular Pathology (ACMG/AMP) guidelines (5). Alternatively, a Bayesian framework consistent with ACMG/AMP guidelines has been proposed (36), in which the likelihood ratios we provide could be used directly and quantitatively to infer variant pathogenicity (in the context of other evidence such as family history, co-segregation, etc.).

A major drawback of our likelihood estimation approach is the limited number of known damaging and tolerated variants currently available. Due to small sample sizes, our current estimate of the score distributions of known damaging and tolerated variants is only an approximation. As more variants in *GDI1* are discovered, assigned clinical significance and added to databases such as ClinVar, this information should be incorporated to more confidently estimate likelihood ratios.

In addition to variant interpretation, variant effect maps can also provide insights into the function of a protein’s structural components. In previous studies, structure-function analysis of GDI1 has been largely focused on the conserved regions common to all members of the *GDI/CHM* superfamily. Our results confirm that variants within the conserved regions forming the Rab binding platform do tend to be the most deleterious. However, certain residues within non-conserved regions exhibited fitness scores that suggested damaging substitutions. These positions may be important for protein folding or stability, or contribute to functional roles of GDI1 not shared by other members of the GDI family. While the MEL region has been the focus for mutational analysis within SCR3A, we found that variants flanking the MEL region, especially within β-strand e3, appeared markedly more deleterious. These findings can be used to guide further mutational analysis of *GDI1*, aimed at discovering the specific functional roles of each of these regions.

Due to the length of the *GDI1* ORF, the coverage of well-measured amino acid substitutions for *GDI1* (19%) was somewhat lower than had been achieved for previous genes studied using this approach (8,9). Nonetheless, precision-recall analysis revealed that, after imputation by machine learning, the variant effect map was able to predict pathogenic variants with greater accuracy than current computational methods alone, and with precision similar to previously studied genes. Thus, using experimental data for a minority of substitutions, we could accurately score variant effects for the majority of amino acid changes.

As genetic testing and exome sequencing continue to be used as diagnostic tools for genetic disorders, it is expected that more patients with novel *GDI1* mutations will be discovered. This map can be used to assist the interpretation of variants immediately upon their discovery, thus accelerating the diagnostic process which is often costly, time-consuming, and stressful for patients and their families. Due to the highly heterogeneous etiology of ID, it is reasonable to expect that response to therapeutic and pharmacological interventions may also vary in accordance with the cause of ID. Unfortunately, therapeutic guidelines rarely differentiate between different forms of ID. Increased rates of etiological diagnoses could improve our understanding of rare forms of ID and aid in the development of more personalized guidelines for management and treatment.

## Conclusions

Here we have presented the first variant effect map for single amino acid substitutions in *GDI1*, and showed that map scores could distinguish presumed-damaging from presumed-tolerated variants with better precision than current computational approaches (including Polyphen2, VARITY, and PROVEAN) at all recall thresholds. Furthermore, our variant effect map recovers known biochemical and structural features of GDI1 and provides insights into structural regions which may be important for GDI1 function.

## Methods

### Strains and Plasmids

The *S. cerevisiae* strain carrying the temperature sensitive Gdi1 allele, TSA64 (*gdi1-1::KanR; his3*Δ*1 leu2*Δ*0 ura3*Δ*0 met15*Δ*0*) (gift from G. Tan, C. Boone and B. Andrews) was used as a host for the *GDI1* variant library. The Gateway destination vector used to express Hs*GDI1*, pHYC-NatMX (CEN/ARS-based, ADH1 promoter, and NatMX marker), was constructed previously (22). The Hs*GDI1* ORF clone (pDONR223-*GDI1*) was obtained from the Human ORFeome v8.1 library (38).

### Construction of *GDI1* variant library by POPcode mutagenesis

POPcode mutagenesis was performed on the *GDI1* ORF as described previously (9): Oligonucleotides of 28-38 bases were designed to target each codon in the open reading frame of *GDI1*, such that the targeted codon is replaced with a NNK-degenerate codon (a mixture of all four nucleotides in the 1st and 2nd codon positions, and a mixture of G and T in the 3rd position). Oligos were annealed to uracilated *GDI1* template, gaps between annealed oligonucleotides were filled using KAPA HiFi Uracil+ DNA polymerase, nicks were sealed using T4 DNA ligase, and the wild type template was degraded using Uracil-DNA-Glycosylase. The variant library was transferred to the yeast expression vector, pHYC-NatMX, by *en masse* Gateway LR reaction (8) followed by transformation into NEB5a competent E. coli cells (New England Biolabs) and selection for ampicillin resistance. Plasmids extracted from a pool of ∼100,000 clones were transformed into the *S. cerevisiae* temperature-sensitive strain TSA64 *en masse* using EZ Kit Yeast Transformation kit (Zymo Research). The entire transformed library was grown in selective media (YPD + clonNAT) for two overnights. All yeast growth was carried out at permissive temperature (25C).

### High-throughput yeast-based complementation

For the pre-selection condition, plasmids were extracted from two 9 ODU samples of yeast culture carrying the variant library (to be used for downstream tiling PCR). For the selective condition, two replicates of 20 ODU of cells were inoculated into 200ml of YPD + clonNAT and grown to full density at restrictive temperature (38°C) with shaking. Plasmids for tiling PCR were extracted from 9 ODU of each culture following competitive growth. In parallel, 2 ODU of TSA64 expressing wild type *GDI1* was inoculated into 20ml of YPD + clonNAT. Wild type pools were grown under the same conditions as the POPcode library and plasmid was extracted from 9 ODU samples to be used as a control for sequencing error during TileSeq.

### Measurement of allele frequencies in pre- and post-selective pools by TileSeq

TileSeq was performed on the plasmids extracted from pre-selective, post-selective, and wild type pools as described previously (8): (i) The *GDI1* ORF was amplified with primers carrying a binding site for Illumina sequencing adaptors; (ii) each amplicon was indexed with an Illumina sequencing adaptor; (iii) paired end sequencing was performed on the tiled amplicons to an average sequencing depth of ∼ 2 million reads. Raw sequencing reads were mapped to the *GDI1* ORF using Bowtie2 (39). A custom Perl script (40) was used to parse the alignment files to count the number of co-occurrences of a codon change in both paired reads. Mutational counts for each tiled region were subsequently normalized by the corresponding sequencing depth, generating a “raw data” file (table S1) where mutational counts are expressed in “reads per million”, i.e. the number of reads normalized to a depth of 1M reads (indicated as “reads/million” below).

### Data processing and fitness score calculation

Processing of raw read count data (available in table S1) was carried out using the “legacy2.R” script (41). This script is derived from the “legacy.R” script from the tileseqMave R package described previously (10), with several modifications to improve filtering and fitness score calculation for variants detected at low frequencies. Read counts for each variant in the wild type control were subtracted from the corresponding read count for variants in each condition in order to account for the detection of variants due to sequencing error. An enrichment ratio (ϕ) was calculated for each variant as the ratio of the normalized read counts after selection to before selection. Since there was less agreement between replicate read counts for variants present at lower frequencies in the pre-selection pool, a pre-filter was applied to remove all variants present in fewer than 200 reads/million in either replicate. The cut-off value of 200 reads/million was chosen in order to maximize the *t*-statistic measure of separation of mean synonymous and pathogenic log ratios (fig. S2a). Additionally, any variants with read counts within 3 standard deviations of zero in the post-selective condition were removed from the data set due to the possibility that they were lost due to a bottleneck effect when sampled from the pre-selective pool. As described previously (8), standard deviation estimates were regularized according to a method for Bayesian regularization described by Baldi and Long (42), which improves confidence estimates for measurements for which few replicates are available (in this case, two). A fitness score (FS_MUT_) was calculated for each variant as ln(ϕ_MUT_/ϕ_STOP_)/ln(ϕ_SYN_/ϕ_STOP_), where ϕ_MUT_ is the enrichment ratio calculated for each variant, ϕ_STOP_ is the median enrichment ratio of all well-measured nonsense variants and ϕ_SYN_ is the median enrichment ratio of all well-measured synonymous variants, such that FS_MUT_ equals zero when ϕ_MUT_ equals ϕ_STOP_ and FS_MUT_ equals one when ϕ_MUT_ equals ϕ_SYN_. Well-measured variants included in the calculation of the medians ϕ_STOP_ and ϕ_SYN_ were those for which enrichment ratios between replicates agreed highly with regularized standard deviation less than 0.3. Because nonsense mutations after residue 400 did not result in complete loss of function (fig. S3), nonsense mutations at amino acid positions greater than 400 were excluded from the ϕ_STOP_ calculation. Fitness scores generated through this pipeline are available in table S2.

### Imputation for missing variant effect map positions and fitness score refinement

Imputation was performed using the variant effect imputation web server (24). The imputation machine learning model was trained on the fitness scores of the experimentally measured variants using the Gradient Boosted Tree (GBT) method. Features of the measured variants used

in the model include mean fitness scores of up to three nearest neighbor variants, standard fitness score error of up to three most similar neighbor variants at the same position, number of neighbors used, PolyPhen-2 score, PROVEAN score, and Blosum100 score. Fitness scores for missing variants were not imputed for positions with fewer than three well-measured variants due to insufficient functional data. Fitness scores of experimentally measured variants were also refined using the weighted average of imputed and measured values (weighting by the inverse-square of estimated standard error in each input value). Output of the imputation pipeline is available in table S3.

### Construction of GDI1 homology model

Human GDI1 (RefSeq: NP_001484.1) residues 1 - 426 were modeled on the crystal structure of RabGDP-dissociation inhibitor in complex with prenylated YPT1 GTPase (PDB: 1UKV) using Swiss-Model ProMod3 Version 1.3.0 (43). The poorly-aligned 21 C-terminal residues were not included in the model.

### Likelihood ratio calculations

Our set of presumed damaging human variants contained Leu92Pro (11), Arg423Pro (15), and Gly237Val (33). Arg423pro is currently annotated as “pathogenic” on ClinVar. Leu92Pro was previously annotated as pathogenic but is currently annotated as having “uncertain significance”, however we believe that d’Adamo et. al (11) provide strong evidence for the deleteriousness of this mutation. Gly237Val was added to ClinVar more recently and is also annotated as having “uncertain significance”, however this variant seemed likely to be deleterious based on familial segregation analysis by Duan et. al (33). We included four additional variants, Tyr39Val, Glu233Ser, Met250Tyr, and Thr248Pro (27), which have not been observed in humans, but which were shown to inhibit GDI1 function in functional assays. The set of presumed tolerated variants consisted of the 46 gnomAD variants from male subjects (hemizygous at the *GDI1* locus), who were presumed to be healthy given that gnomAD excludes subjects with early childhood disease. Normal distributions were fitted to the histograms of the fitness scores of presumed damaging and tolerated variants by maximum likelihood parameter estimation in order to obtain estimated probability density functions for pathogenic/disease and benign variants (*p*_*D*_ and *p*_*B*_ respectively). The normal distributions used are shown in fig. 4a (but scaled such that the area under each curve equals 1 for likelihood ratio calculations). The damaging:tolerated likelihood ratio for a variant with fitness score, *f*, was calculated as the ratio of the estimated probability density functions evaluated at *f*: Λ(*D*: *T* | *f*) = *p*_*D*_(*f*)/*p*_*T*_(*f*). This likelihood ratio can be used together with prior beliefs about a variants’ pathogenicity to calculate the odds that a variant is damaging, *O*(*D*: *T* | *f*),using the Odds form of Bayes’ rule:

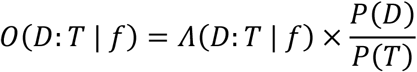

where, Λ(*D*: *T* | *f*) is the likelihood ratio, *P*(*D*) is the prior probability that the variant is damaging, and *P*(*T*) is the prior probability that the variant is tolerated such that *P*(*T*) = 1 ™ *P*(*D*).

## Declarations

### Ethics approval and consent to participate

Not applicable

### Consent for publication

Not applicable

### Availability of data and materials

All data generated or analyzed during this study are included in this published article and its supplementary information files.

### Authors’ contributions

SS and FPR conceived the project. SS established the assay. MV, SS, IF, and JK performed mutagenesis and selection. MG performed sequencing. RAS performed primary data analyses with contributions from SS and JW. RAS performed all downstream analyses of map scores, including analysis of sequence-structure-function relationships. YW developed the machine learning imputation pipeline with contributions from JW. RAS and FPR wrote the manuscript. All authors read and approved the manuscript.

## Acknowledgments

We thank Guihong Tan, Charles Boone and Brenda Andrews for generously providing the TSA64 *S. cerevisiae* strain.

## Supplemental Information

### Supplemental Figures and Legends

**Figure S1:**
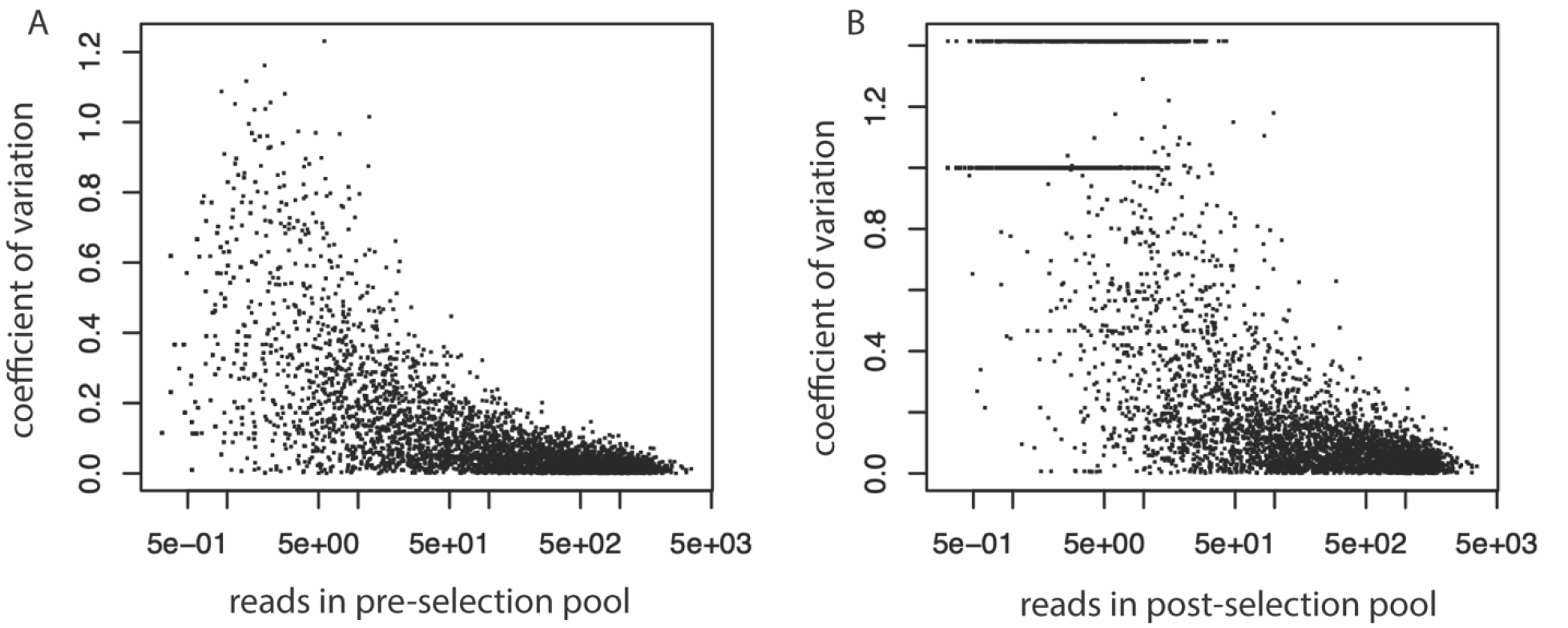
Variants present at low frequencies in complementation screen show poorer agreement between replicates. a) Coefficient of variation between two read count replicates for all detected variants in the pre-selection pool versus frequency in the pre-selection pool (as measured by mean read count of the two replicates). b) Coefficient of variation between two read count replicates for all detected variants in the post-selection pool versus frequency in the post-selection pool (as measured by mean read count of the two replicates).

**Figure S2:**
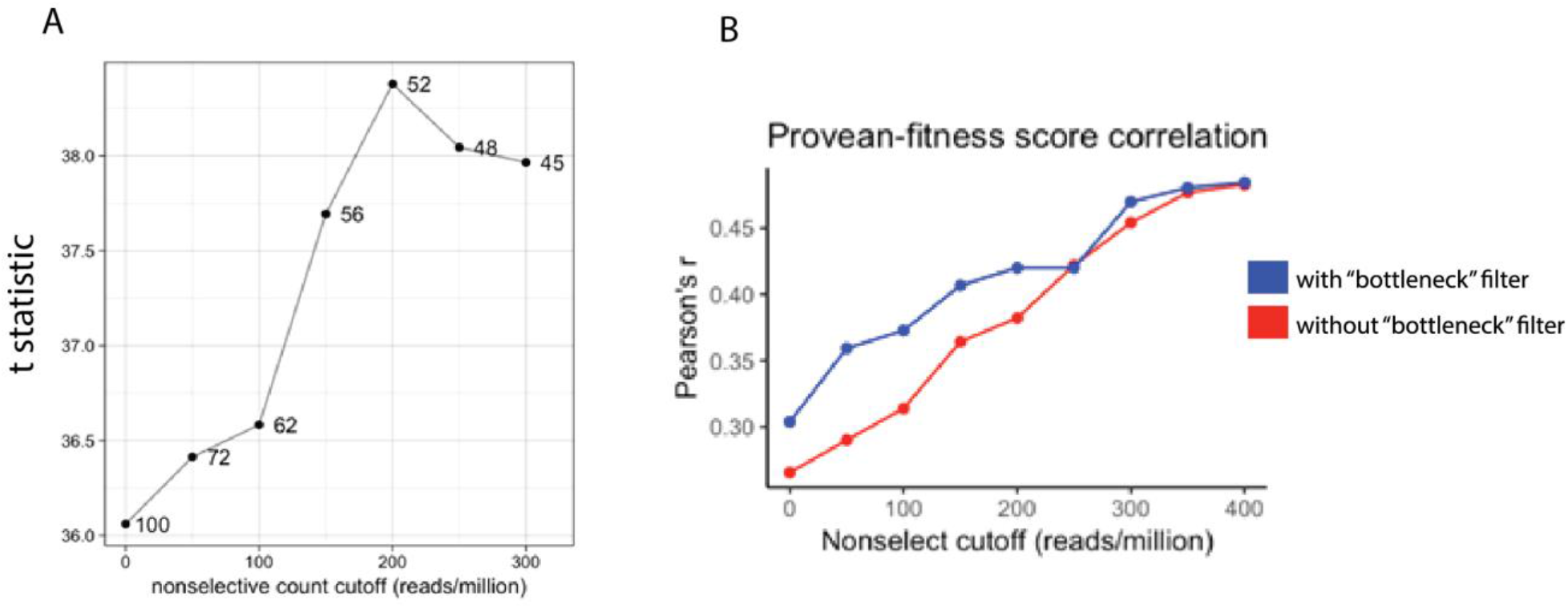
Filtering out variants present at low frequencies in the pre-selection pool improves metrics of fitness measurement accuracy. a) Multiple read count cut-offs were tested wherein read counts in the pre-selection pool were filtered to include only high-frequency variants (present at frequencies greater than the cut-off value). For each cut-off value tested, a two-sample t-statistic was calculated to evaluate the separation of fold changes between nonsense variants and synonymous variants. A cut-off value of 200 reads/million maximized the separation of synonymous and nonsense variants. b) Correlation between PROVEAN scores and our fitness scores increase as variants are filtered for higher frequency in the pre-selection variant pool. For each read count cutoff, the correlation (Pearson’s R) between our calculated fitness score (prior to imputation) and PROVEAN scores for all missense variants was calculated.

**Figure S3:**
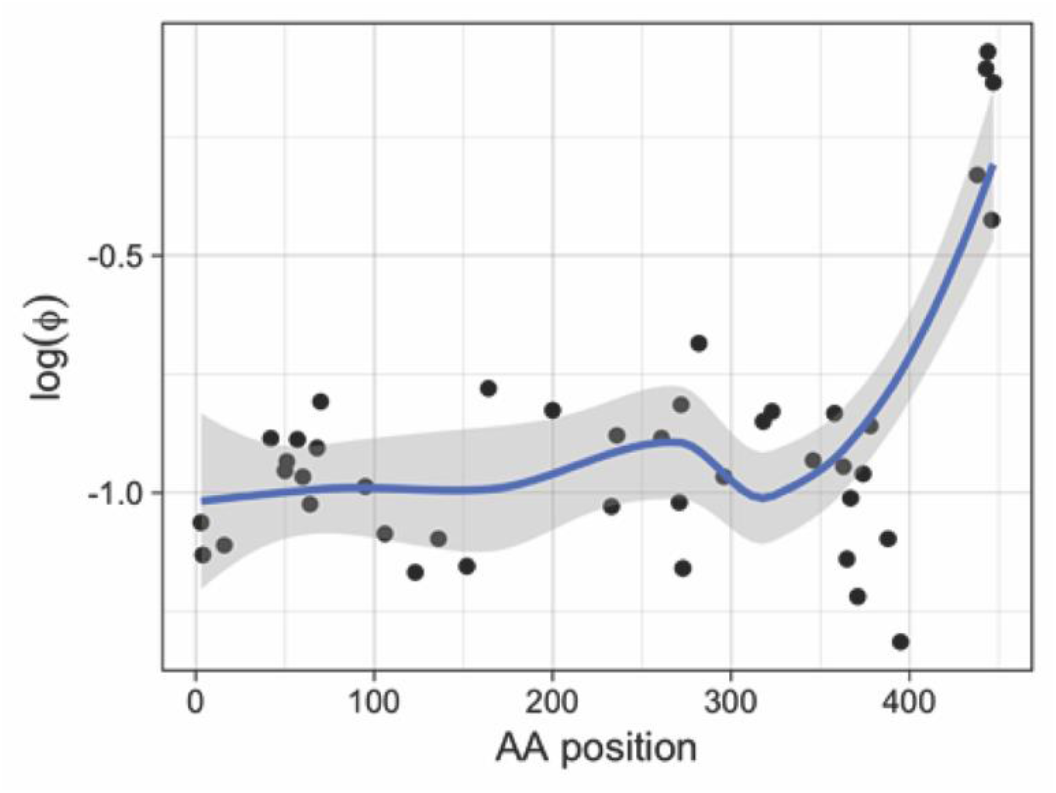
Fold changes (pre-selection/post-selection) of all measured nonsense mutations in *GDI1*. Nonsense mutations after amino acid position 400 lead to less severe loss of complementation.

**Figure S4:**
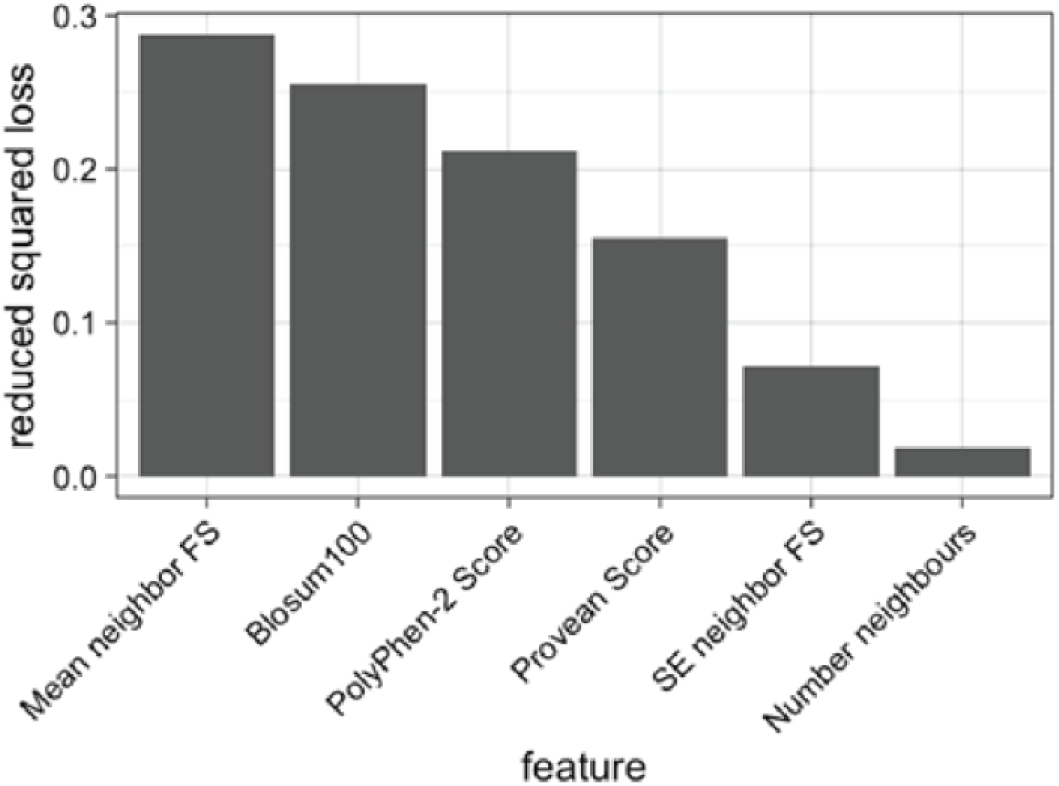
Feature importance for gradient boosted trees imputation model. Mean neighbor FS: the mean fitness scores of the 3 most similar amino acids at the same residue position. SE neighbor FS: Standard error of the fitness scores of the 3 most similar amino acids at the same residue position. Number neighbors: Number of variants measured at the same amino acid position

### Descriptions of Supplementary Tables

**Table S1: Table of raw yeast complementation data**

Table of unfiltered *GDI1* variant frequencies in pre- and post-selection deep sequencing pools. Variants counts are presented in reads/million.

**Table S2: Fitness score table**

Table of fitness score data calculated for all well-measured *GDI1* variants.

**Table S3: Imputed scores**

Table of all fitness scores including computationally imputed scores for amino acid substitutions not measured experimentally.

## References

1. Karczewski KJ, Francioli LC, Tiao G, Cummings BB, Alföldi J, Wang Q, et al. The mutational constraint spectrum quantified from variation in 141,456 humans. Nature. 2020;

2. Starita LM, Ahituv N, Dunham MJ, Kitzman JO, Roth FP, Seelig G, et al. Variant Interpretation: Functional Assays to the Rescue. American Journal of Human Genetics. 2017.

3. Landrum MJ, Lee JM, Benson M, Brown G, Chao C, Chitipiralla S, et al. ClinVar: Public archive of interpretations of clinically relevant variants. Nucleic Acids Res. 2016;

4. Wright CF, West B, Tuke M, Jones SE, Patel K, Laver TW, et al. Assessing the Pathogenicity, Penetrance, and Expressivity of Putative Disease-Causing Variants in a Population Setting. Am J Hum Genet. 2019;

5. Richards S, Aziz N, Bale S, Bick D, Das S, Gastier-Foster J, et al. Standards and guidelines for the interpretation of sequence variants: A joint consensus recommendation of the American College of Medical Genetics and Genomics and the Association for Molecular Pathology. Genet Med. 2015;

6. Fowler DM, Araya CL, Fleishman SJ, Kellogg EH, Stephany JJ, Baker D, et al. High-resolution mapping of protein sequence-function relationships. Nat Methods. 2010;

7. Fowler DM, Fields S. Deep mutational scanning: A new style of protein science. Nature Methods. 2014.

8. Weile J, Sun S, Cote AG, Knapp J, Verby M, Mellor JC, et al. A framework for exhaustively mapping functional missense variants. Mol Syst Biol. 2017;

9. Sun S, Weile J, Verby M, Wu Y, Wang Y, Cote AG, et al. A proactive genotype-to-patient-phenotype map for cystathionine beta-synthase. Genome Med. 2020;

10. Weile J, Kishore N, Sun S, Maaieh R, Verby M, Li R, et al. Shifting landscapes of human MTHFR missense-variant effects. Am J Hum Genet. 2021;

11. D’Adamo P, Menegon A, Nigro C Lo, Grasso M, Gulisano M, Tamanini F, et al. Mutations in GDI1 are responsible for X-linked non-specific mental retardation. Nat Genet. 1998;

12. D’Adamo P, Welzl H, Papadimitriou S, Di Barletta MR, Tiveron C, Tatangelo L, et al. Deletion of the mental retardation gene Gdi1 impairs associative memory and alters social behavior in mice. Hum Mol Genet. 2002;

13. Potokar M, Jorgačevski J, Lacovich V, Kreft M, Vardjan N, Bianchi V, et al. Impaired αGDI Function in the X-Linked Intellectual Disability: The Impact on Astroglia Vesicle Dynamics. Mol Neurobiol. 2017;

14. Goody RS, Rak A, Alexandrov K. The structural and mechanistic basis for recycling of Rab proteins between membrane compartments. Cellular and Molecular Life Sciences. 2005.

15. Bienvenu T, Des Portes V, Saint Martin A, McDonell N, Billuart P, Carrié A, et al. Non-specific X-linked semidominant mental retardation by mutations in a Rab GDP-dissociation inhibitor. Hum Mol Genet. 1998;

16. Strobl-Wildemann G, Kalscheuer VM, Hu H, Wrogemann K, Ropers HH, Tzschach A. Novel GDI1 mutation in a large family with nonsyndromic X-linked intellectual disability. Am J Med Genet Part A. 2011;

17. Stessman HAF, Xiong B, Coe BP, Wang T, Hoekzema K, Fenckova M, et al. Targeted sequencing identifies 91 neurodevelopmental-disorder risk genes with autism and developmental-disability biases. Nat Genet. 2017;

18. Kvarnung M, Nordgren A. Intellectual disability & rare disorders: A diagnostic challenge. In: Advances in Experimental Medicine and Biology. 2017.

19. Bélanger SA, Caron J. Evaluation of the child with global developmental delay and intellectual disability. Paediatr Child Heal. 2018;

20. Monroe GR, Frederix GW, Savelberg SMC, De Vries TI, Duran KJ, Van Der Smagt JJ, et al. Effectiveness of whole-exome sequencing and costs of the traditional diagnostic trajectory in children with intellectual disability. Genet Med. 2016;

21. Thevenon J, Duffourd Y, Masurel-Paulet A, Lefebvre M, Feillet F, El Chehadeh-Djebbar S, et al. Diagnostic odyssey in severe neurodevelopmental disorders: Toward clinical whole-exome sequencing as a first-line diagnostic test. Clin Genet. 2016;

22. Sun S, Yang F, Tan G, Costanzo M, Oughtred R, Hirschman J, et al. An extended set of yeast-based functional assays accurately identifies human disease mutations. Genome Res. 2016;

23. Choi Y, Chan AP. PROVEAN web server: A tool to predict the functional effect of amino acid substitutions and indels. Bioinformatics. 2015;

24. Wu Y, Weile J, Cote AG, Sun S, Knapp J, Verby M, et al. A web application and service for imputing and visualizing missense variant effect maps. Bioinformatics. 2019;

25. Henikoff S, Henikoff JG. Amino acid substitution matrices from protein blocks. Proc Natl Acad Sci U S A. 1992;

26. Adzhubei IA, Schmidt S, Peshkin L, Ramensky VE, Gerasimova A, Bork P, et al. A method and server for predicting damaging missense mutations. Nature Methods. 2010.

27. Schalk I, Zeng K, Wu SK, Stura EA, Matteson J, Huang M, et al. Structure and mutational analysis of Rab GDP-dissociation inhibitor. Nature. 1996;

28. Rak A, Pylypenko O, Durek T, Watzke A, Kushnir S, Brunsveld L, et al. Structure of Rab GDP-Dissociation Inhibitor in Complex with Prenylated YPT1 GTPase. Science (80-). 2003;

29. Luan P, Heine A, Zeng K, Moyer B, Greasely SE, Kuhn P, et al. A new functional domain of guanine nucleotide dissociation inhibitor (alpha-GDI) involved in Rab recycling. Traffic. 2000;

30. An Y, Shao Y, Alory C, Matteson J, Sakisaka T, Chen W, et al. Geranylgeranyl switching regulates GDI-Rab GTPase recycling. Structure. 2003;

31. Patel DR, Apple R, Kanungo S, Akkal A. Intellectual disability: Definitions, evaluation and principles of treatment. Pediatric Medicine. 2018.

32. Hamel BCJ, Kremer H, Wesby-van Swaay E, Van Den Helm B, Smits APT, Oostra BA, et al. A gene for nonspecific X-linked mental retardation (MRX41) is located in the distal segment of Xq28. Am J Med Genet. 1996;

33. Duan Y, Lin S, Xie L, Zheng K, Chen S, Song H, et al. Exome sequencing identifies a novel mutation of the GDI1 gene in a Chinese non-syndromic X-linked intellectual disability family. Genet Mol Biol. 2017;

34. Adzhubei I, Jordan DM, Sunyaev SR. Predicting functional effect of human missense mutations using PolyPhen-2. Curr Protoc Hum Genet. 2013;

35. Wu Y, Li R, Sun S, Weile J, Roth FP. Improved pathogenicity prediction for rare human missense variants. Am J Hum Genet. 2021;

36. Tavtigian S V., Greenblatt MS, Harrison SM, Nussbaum RL, Prabhu SA, Boucher KM, et al. Modeling the ACMG/AMP variant classification guidelines as a Bayesian classification framework. Genet Med. 2018;

37. NHLBI GO Exome Sequencing Project (ESP). Exome Variant Server. NHLBI. 2018.

38. Yang X, Boehm JS, Yang X, Salehi-Ashtiani K, Hao T, Shen Y, et al. A public genome-scale lentiviral expression library of human ORFs. Nat Methods. 2011;

39. Langmead B, Salzberg SL. Fast gapped-read alignment with Bowtie 2. Nat Methods. 2012;

40. TileSeq package [Internet]. Available from: https://github.com/rothlab/tileseq_package

41. Weile J, Silverstein R. Github/tileseqMave/legacy2.R [Internet]. Available from: https://github.com/RachelSilverstein/tileseqMave/blob/master/R/legacy2.R

42. Baldi P, Long AD. A Bayesian framework for the analysis of microarray expression data: Regularized t-test and statistical inferences of gene changes. Bioinformatics. 2001;

43. Studer G, Tauriello G, Bienert S, Biasini M, Johner N, Schwede T. ProMod3 - A versatile homology modelling toolbox. PLoS Comput Biol. 2021;

44. McGuffin LJ, Bryson K, Jones DT. The PSIPRED protein structure prediction server. Bioinformatics. 2000;

